# EDGE2: advancing the prioritisation of threatened evolutionary history for conservation action

**DOI:** 10.1101/2022.05.17.492313

**Authors:** Rikki Gumbs, Claudia L. Gray, Monika Böhm, Ian J. Burfield, Olivia R. Couchman, Daniel P. Faith, Félix Forest, Michael Hoffmann, Nick J. B. Isaac, Walter Jetz, Georgina M. Mace, Arne O. Mooers, Kamran Safi, Oenone Scott, Mike Steel, Caroline M. Tucker, William D. Pearse, Nisha R. Owen, James Rosindell

## Abstract

The global biodiversity crisis threatens the natural world and its capacity to provide benefits to humans into the future. The conservation of evolutionary history, captured by the measure phylogenetic diversity (PD), is linked to the maintenance of these benefits and future options. The Evolutionarily Distinct and Globally Endangered (EDGE) metric has, since 2007, been used to identify species for conservation action that embody large amounts of threatened evolutionary history. In 2017, we convened a workshop to update the EDGE metric to incorporate advances in the field of phylogenetically-informed conservation. Building on that workshop, we devised the metric ‘EDGE2’, which we present here. EDGE2 uses a modular, tiered approach to provide priority rankings—and associated measures of uncertainty in both phylogenetic and extinction risk data—for all species in a clade. EDGE2 takes into account the extinction risk of closely-related species to better reflect the contribution a species is expected to make to overall PD in the future. We applied EDGE2 to the world’s mammals to identify an updated list of priority EDGE species and compare the results with the original EDGE approach. Despite similarity in the priority lists produced between EDGE and EDGE2, EDGE2 places greater priority on species with fewer close relatives on the Tree of Life. As we approach a crossroads for global biodiversity policy, EDGE2 exemplifies how academic and applied conservation biologists can collaborate to guide effective priority-setting to conserve the most irreplaceable components of biodiversity upon which humanity depends.

## Introduction

### Why conserve evolutionary history?

Global declines in biodiversity imperil nature’s capacity to provide benefits critical to maintaining the quality of human life now and into the future [1]. There are, however, different currencies by which biodiversity is measured and prioritised, one of which considers the evolutionary history contained in a set of taxa. The biodiversity measure phylogenetic diversity (PD [2]), calculated by summing the phylogenetic branch lengths spanning a set of taxa, links evolutionary history to the conservation of feature diversity (the different evolutionary features of species), and so to future options for humanity (or ‘biodiversity option value’ [2,3]). Therefore, by preserving PD, and its associated diversity of features, we can maintain the benefits and future options these features contribute to humanity [4–7].

Conservation initiatives that incorporate evolutionary history have the potential to capture other desirable components of biodiversity (e.g. ‘functional’ trait diversity [8–11]), and the Tree of Life itself reflects a fundamental component of biodiversity with its own intrinsic value [12–15]. The importance of halting the loss of evolutionary history has been recognised by the Members of the International Union for Conservation of Nature (IUCN), who adopted Resolution WCC-2012-Res-019-EN, which calls for more “conservation initiatives that target species, especially those of high evolutionary significance” [16]. In light of this, the IUCN has now established a Phylogenetic Diversity Task Force to provide expertise on the inclusion of PD in conservation strategies for practitioners, decision-makers, and the public [17]. Further, the Intergovernmental Science-policy Platform for Biodiversity and Ecosystem Services (IPBES) has adopted PD as an indicator of the overall capacity of biodiversity to support a good quality of life into the future (the ‘maintenance of options’ [5]). This indicator, along with an index tracking the conservation of the most evolutionarily unique and threatened species (the ‘EDGE Index’), have also been included as indicators for the Convention on Biological Diversity’s draft post-2020 Global Biodiversity Framework [18].

### The history of the EDGE metric

In 2007, the Zoological Society of London (ZSL) established the ‘EDGE’ metric as a method for identifying species that should be prioritised for the conservation of threatened evolutionary history. The approach relied on combining a measure of Evolutionary Distinctiveness (ED) with values for species’ risk of extinction (Global Endangerment - GE) to calculate ‘EDGE scores’ and thereby generating priority rankings [19], or ‘EDGE Lists’.

ED assigns each species a ‘fair proportion’ of the total PD of the phylogeny: each species that is descended from a given phylogenetic branch receives an equal proportion of the length of that branch’s PD [20]. Species with long ancestral branches that are shared with relatively few other species are therefore responsible for greater amounts of PD than species with short ancestral branches that are shared with numerous other species. The ED of species *i* can be written as:

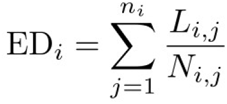

Here *L_i_*_,1_ gives the terminal branch length of species *i*, *L_i,j_* for 2 ≤ *j* ≤ *n_i_* gives the length of all internal branches that are ancestral to species *i* and *N_i,j_* gives the total number of descendants of each of these same branches.

GE utilised weightings of extinction risk derived from the categories of the IUCN Red List of Threatened Species that are already widely used to produce Red List Indices [21]. Following Isaac et al. [19], the GE of species *i* (*GE_i_*) was given as 0 where species *i* is ‘Least Concern’ (LC), 1 where it is ‘Near Threatened’ (NT), 2 where it is ‘Vulnerable’ (VU), 3 where it is ‘Endangered’ (EN), and 4 where it is ‘Critically Endangered’ (CR). Hereafter, we refer collectively to these as “data-sufficient” categories. In the original metric, Extinct in the Wild species were not included. The EDGE score for any species *i* was then calculated as:

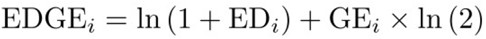

Following Isaac et al. [19], species in threatened IUCN Red List categories of VU, EN, or CR (collectively referred to as “threatened”) with above-median ED for their clade were identified as priority ‘EDGE Species’, with particular conservation attention given to the highest-ranking 100, 50 or 25 species in particular clades [22,23].

These priority EDGE Lists have been generated for mammals [19,24], amphibians [25], birds [26], corals [27], reptiles [22], gymnosperms [28] and sharks and rays [29]. EDGE Lists have informed direct conservation action for many of the EDGE Species highlighted, and have served as the basis for conservation efforts of ZSL’s own EDGE of Existence programme. The EDGE programme has carried out over 120 conservation projects worldwide on priority EDGE Species, and regularly updated EDGE Lists are hosted at www.edgeofexistence.org.

ED scores have often been utilised as a measure of species distinctiveness [29–32], and EDGE Species are increasingly recognised as being of global conservation importance [16,33,34]. EDGE data underpins initial estimations by IPBES for their PD indicator [5,35,36], which approximates expected loss of PD [37]. EDGE data also informs conservation grant mechanisms, such as the IUCN Species Survival Commission (SSC) EDGE Internal Grant [38], which funds Red Listing and action planning for evolutionarily distinct species and lineages, and the conservation needs assessments utilised by Amphibian Ark’s conservation grants programme [39].

### Revisiting the EDGE metric

The success in uptake of PD in research, policy, and applied conservation has occurred alongside great advances across the field of phylogenetically-informed conservation prioritisation [26,40–44]. The original EDGE metric, whilst having strong theoretical underpinnings, is an heuristic measure designed to facilitate swift judgments on prioritising species for conservation—according to the information derived from two elements: the ED (irreplaceability), and the GE (vulnerability) of species—which result in tangible conservation action. The advances in the field of PD conservation have opened up new avenues to produce a mechanistic EDGE metric that also embodies these desirable properties. Relevant developments included the quantification of extinction risk [42,45], the incorporation of uncertainty in both phylogeny [22,26,46] and extinction risk [47,48], and the concept of phylogenetic complementarity between species [40,41,49]. Each of these developments indicates a potential weakness in the original EDGE metric and a corresponding opportunity to improve its performance.

We convened a workshop in April 2017 to assess whether the EDGE metric should be formally updated in light of a decade of research advances since 2007. Given that the expected outcome of this workshop would potentially alter species priorities for conservation, it was necessary to carefully assess and justify any changes made. We brought together a wide variety of experts, and required a clear majority agreement on all decisions made so as to produce a credible, comprehensive, and robust update. Here, we introduce the resulting new EDGE metric, hereafter referred to as ‘EDGE2’. We characterise the key advantages of EDGE2 as a tool to conserve threatened evolutionary history and, finally, we provide guidance for interpreting EDGE2, illustrated with an application to the world’s mammals.

### Introducing EDGE2

The EDGE2 metric incorporates earlier probabilistic approaches to phylogenetically-informed conservation prioritisation [40,41,50,51] to measure the avertable loss of phylogenetic diversity through the conservation of individual species. We frame ‘EDGE2’ in terms of the product between an ‘ED2’ component (the irreplaceability of a species) and a ‘GE2’ component (the vulnerability of a species) of species *i*:

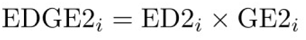

This approach follows earlier formulations to calculate species-specific expected losses of evolutionary history [52] to provide a measure of avertable expected PD loss [37,40]. The approach emphasises modularity, a key design principle of EDGE2, which allows future research developments to fit into either component separately.

### GE2 - Global Endangerment

The modular nature of EDGE2 means that GE2 remains a distinct component of the metric, as with original GE. In the original formulation of EDGE, the metric dictates that, should the extinction risk of all species be equal, the most evolutionarily distinct species will be prioritised [19]. However, given that extinction risk varies across species, the original formulation of GE was designed to weight the ED of species so that high ED species would not be ignored in the presence of the many species that are more threatened, but less distinct.

We continue to regard GE2*_i_* as a global endangerment weighting, now between 0 and 1, for species *i* relative to other species. For simplicity in this manuscript, we set GE2*_i_* = *p_i_*, where *p_i_* gives the probability of extinction for species *i* at some given future time. However, in the spirit of EDGE2 modularity, other choices could be used in future. For example, if we are evaluating the potential impacts of particular conservation actions, GE2*_i_* may reflect the likely reduction in probability of extinction from a particular action [41,43,53]. Choosing GE2*_i_* = *p_i_* means we take a particular conservation action to mean species *i* is protected from extinction(following [40]; theorem 4.1 (i)).

#### Probability of extinction

The perfect extinction risk data would be probabilities of extinction for all species at all future points in time with no error. However, the best available extinction risk data for most species are the discrete categories of the IUCN Red List [54], which are themselves not available for a large proportion of biodiversity (e.g. only ca. 10% of plants and 0.2% of fungi have been formally assessed [55]). There are no universally accepted corresponding probabilities of extinction for the Red List categories [21,56–58]. A range of quantitative analyses enable species to be assessed under criterion E of the Red List [54], which does directly correspond to a probability of extinction. However, criterion E is very rarely used [59], and we are a long way from having quantitatively calculated species-specific extinction probabilities for entire clades. General probabilities of extinction for each Red List category have been derived from the thresholds of criterion E and scaled to various timeframes [42,52,60], although this has been criticised [56,57,59,61].

It is, however, reasonable to assume that extinction risk is higher amongst Critically Endangered species than for those in Endangered and Vulnerable categories, at least on average. Indeed, Butchart et al. [21]indicate that extinction risk may increase by as much as an order of magnitude for every step-wise increase in Red List category. We chose to map GE2 to Red List categories in a way that preserves the same intervals between Red List categories as the original GE index, where each step-wise increase in Red List category results in a doubling of extinction risk [19]. This followed the principle of retaining components from the original EDGE unless we have a compelling reason to do otherwise. It also means that we only needed to establish a probability of extinction for Critically Endangered species and the rest would follow.

At our workshop, we found no compelling argument as to what time horizon would be most appropriate for calculating probabilities of extinction (e.g. those of Mooers et al. [42]). Delegates could only agree to dismiss time horizons that were either too soon (e.g. 10 years) or too distant (e.g. 500 years) to be relevant for stimulating action. There are naturally subjective elements to any choice; it is tempting to advocate for time horizons farther into the future based on an intuition that conservation efforts are for long-term benefits and should therefore be protected from potential short-sighted decision-making. However, such arguments may not hold up to further scrutiny. Extinction risk information for species can be expected to be updated at the scale of years rather than decades; IUCN Red List assessments of extinction risk are deemed out-dated after 10 years [54]. Although timely reassessments of species can be difficult to achieve [62,63], we still see this as reasonable justification for choosing a time horizon on the order of decades rather than centuries.

We therefore opted to tie the doubling feature of GE’s extinction risk to the 50-year time horizon specified in Mooers et al. [42] (a study notable for examining the impact of time-horizon and *p* definition on conservation prioritisation), with Critically Endangered (CR) mapped to an extinction risk weighting of 0.97. Endangered (EN) species therefore receive a weighting of 0.485, Vulnerable (VU) = 0.2425, Near Threatened (NT) = 0.12125, and Least Concern (LC) = 0.060625.

Our choice of an absolute extinction risk value for CR species tied to the 50-year time horizon, combined with the principle of doubling extinction risk with each increase in Red List category from the original EDGE metric, provides a set of values to capture the original properties of the GE component. That is, the extinction risk values are of high absolute value and risk is halved with every down listing, consistent with the original EDGE formulation [19,25]. At the workshop we considered many other alternatives (e.g.[42]), but no single approach received enough support to justify a further departure from the original EDGE approach. Thus, the concept of GE2 represents a clear conceptual advance in some respects, allowing for the incorporation of probabilities of extinction, while retaining the behaviour of original GE scores in other respects where current knowledge is not enough to make an unequivocal improvement.

#### Uncertainty in probability of extinction

We developed a new approach to incorporate uncertainty in probability of extinction and enable the inclusion of Data Deficient (DD) and Not Evaluated (NE) species, the inclusion of which has been demonstrated to reduce bias in estimates of phylogenetically-informed prioritisations [48]. Our approach is based on the idea that if we rank species by their true probability of extinction, the probabilities will change smoothly reflecting biological processes, and not jump as we move between discrete, arbitrary, human-inferred Red List categories (Fig 1).

**Fig 1:**
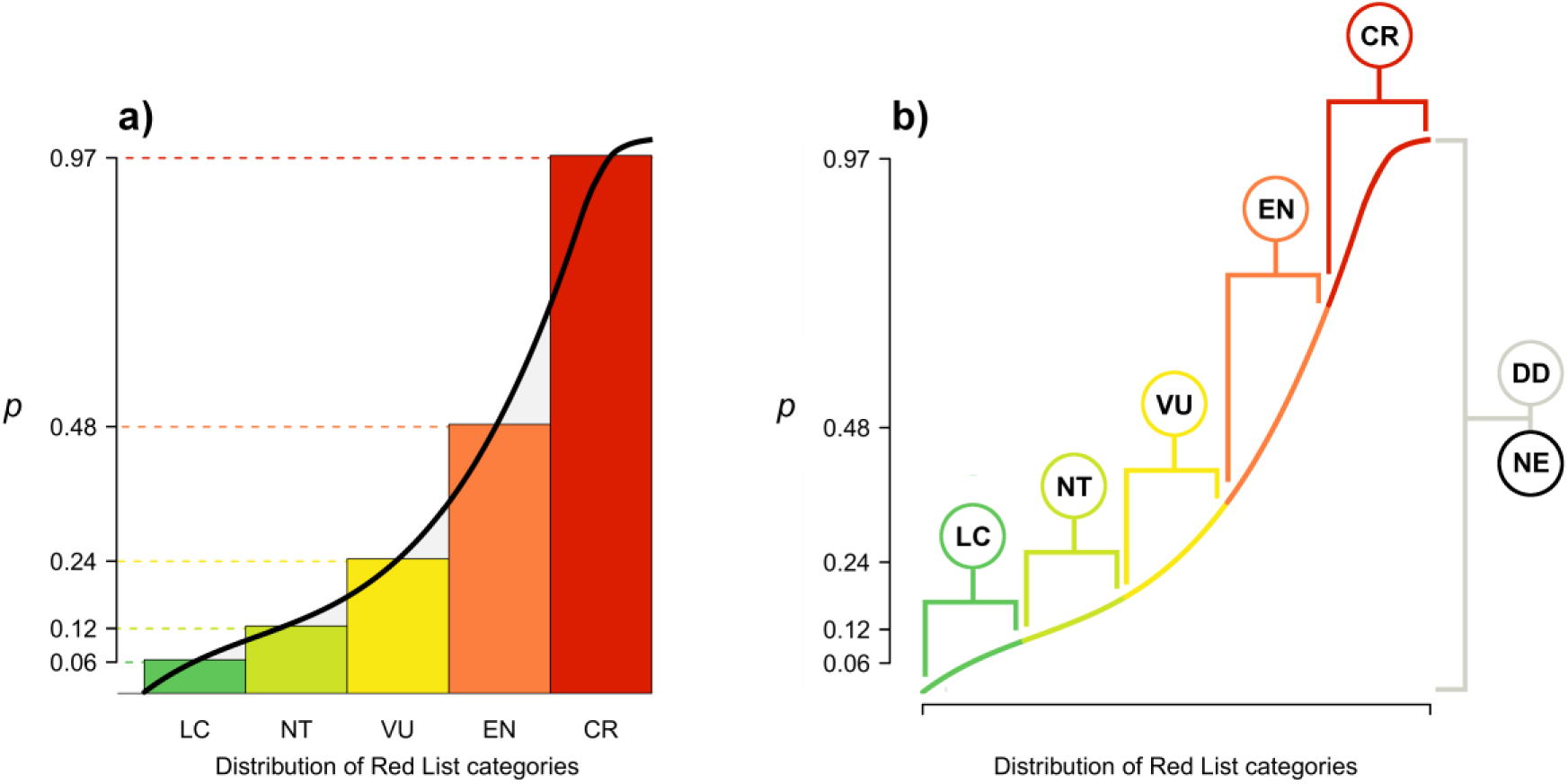
generating a distribution of probabilities of extinction (*p*) from discrete IUCN Red List categories. To generate a continuous distribution of values from which to draw *p* (a), we created a rank plot of ranked extinction risk (x-axis) assigned to the designated median *p* for each IUCN Red List category (y-axis; see ‘probability of extinction’ section). We fitted a quartic curve through the five median values for each Red List category and bounded the resulting curve to return values between 0.0001 and 0.9999 (not wishing to ever return a 0 or 1 extreme) - see methods. From this curve, we could extract a set of values associated with each Red List category (b), the median of which was equivalent to the pre-defined *p* for each category (CR = 0.97; EN = 0.485; VU = 0.2425; NT = 0.12125; LC = 0.060625). The *p* for Data Deficient (DD) and unassessed (Not Evaluated; NE) species is drawn from the entire distribution to capture uncertainty around their extinction risk.

We can therefore reconstruct a smooth curve based on realistic constraints: (1) *p* bounded by a maximum of 0.9999 (almost certain to go extinct) at one extreme and 0.0001 (safe) at the other extreme; (2) *p* must be increasing on account of being ranked by increasing extinction risk; and (3) the median *p* drawn from the portion of distribution corresponding to a Red List category must equal the chosen *p* for that category - the median of all possible *p* draws for a CR species. should remain equal to 0.97 after the process (Fig 1). The portion of the distribution assigned to each category is equal in size, though this could also vary in size in proportion to the observed distribution of Red List categories for a taxonomic group. We elected to use equal bands for each category to allow the generated distribution to be more widely applicable and consistent across multiple taxa, for which the distribution of Red List categories differs. For each species, *p* is drawn for each species from the rank plot by selecting at random a value on the x-axis that falls within the range for the corresponding Red List category (Fig 1b).

For unassessed NE and DD species, *p* is drawn from the entire distribution completely at random under the assumption that, although we do not know where NE and DD species will be located along the curve, it must be at some point between 0 and 1, as with other extant species. Thus we generate highly uncertain scores for NE and DD species that have a median value equivalent to elevated extinction risk (VU) when taken over a large number of iterations [64,65]. For species listed as Possibly Extinct (CR(PE)), Possibly Extinct in the Wild (CR(PEW)) and Extinct in the Wild (EW), *p* is drawn from the corresponding CR values, and all Extinct (EX) species have *p* = 1 by definition. The treatment of PE, PEW, and EW as distinct from EX, with *p* equal to CR, deviates from the Red List Index approach [21]; we took this approach to reflect the fact that PE/PEW/EW species, unlike EX species, have the potential for recovery in the wild.

We draw *p* for all species again in each iteration of calculating EDGE2 and can therefore capture uncertainty in the final results of ED2 and GE2 (= *p*) while retaining the condition that the median GE2 score for each species aligns with the score corresponding to its Red List category. Other sources of uncertainty, for example in the Red List categorisation for the species themselves, and in the treatment of CR(PE), CR(PEW), and EW species, are subjects for future research. For clades where only a single well-supported consensus phylogenetic tree containing all recognised species is available, ED2 must still be calculated a large number of times to adequately capture the uncertainty in *p* (100 < n ≤ 1000 to mirror approaches to capture phylogenetic uncertainty [24,26,46], discussed later) .

#### ED2

One attractive element of the original EDGE formulation is that individual species were assigned a measure of distinctiveness—or irreplaceability—based on their phylogenetic position and relationships: their Evolutionary Distinctiveness (ED [19]). We retain this critical component in EDGE2 whilst introducing PD complementarity. This means that when calculating ED for a species, we consider the extinction risk of its close relatives to better reflect the expected contribution of the species to the future phylogenetic diversity (Fig 2) [40,41,66]. For example, consider we a species that is Critically Endangered, but where the sister species (i.e. they share the same most recent internal phylogenetic branch) is Least Concern. Under the original ED formulation, the contribution of the shared ancestral branch to the ED of the Critically Endangered species is unaffected by the extinction risk of the sister species. However, when we incorporate PD complementarity, the contribution of the shared internal branch to the ED score of the Critically Endangered species is affected by the extinction risk of the sister species. As the sister species is Least Concern, and unlikely to become extinct, the shared internal branch is quite safe, and thus contributes less to the ED score than if the sister species was also threatened with extinction. Had the sister species been Critically Endangered, the internal branch would have been at much greater risk and thus contribute more to the ED score of our focal species. Thus, in our idealised scenario, ED2 measures the expected phylogenetic contribution of a species based on the probability that it will become the sole extant descendant of any of its ancestral phylogenetic branches (i.e. the probability that all other descendants become extinct) at a future point in time.

**Fig 2:**
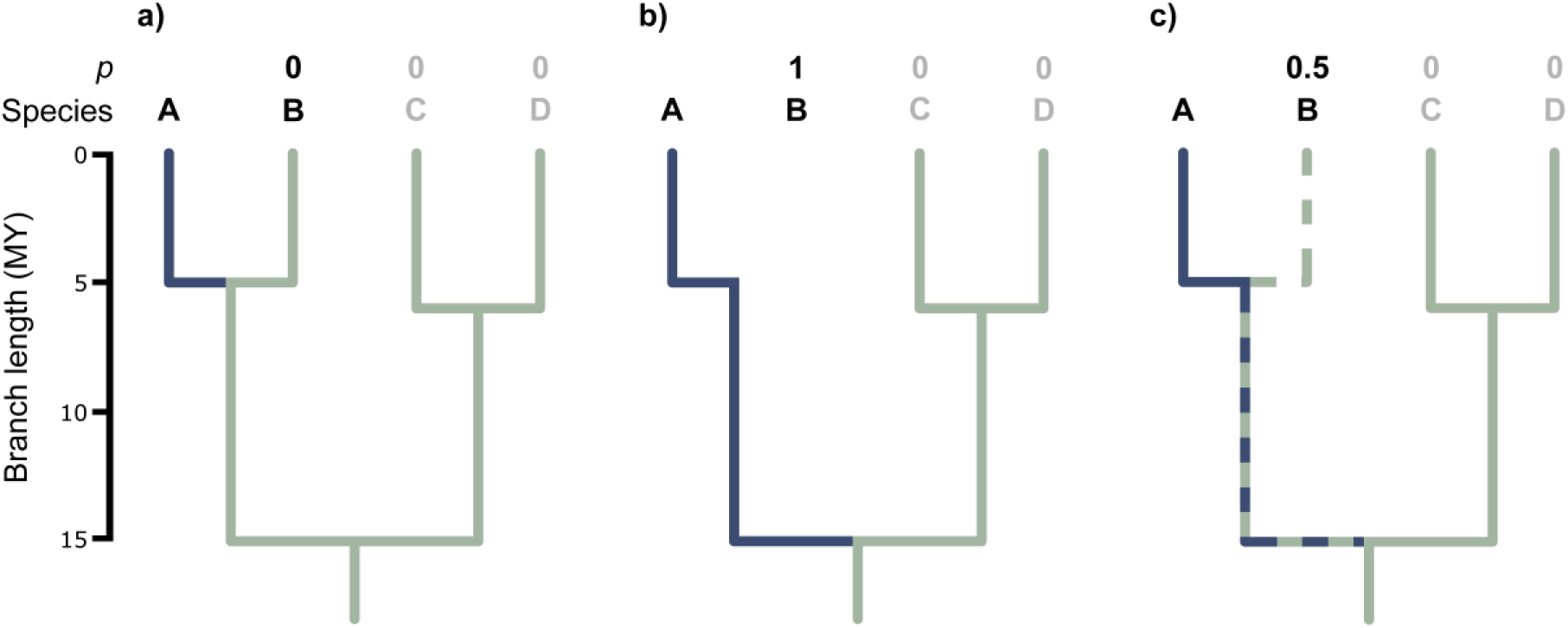
Incorporating PD complementarity into ED2 calculations under different extinction scenarios. When the extinction risk (*p*; see ED2 equation in ‘EDGE2’ section) of all species is 0, as in panel (a), the ED2 of species A is limited to the length of its terminal branch (blue branch), as there is zero probability of species B becoming extinct, and therefore zero probability their shared internal branch will contribute to species A’s terminal branch length in the future. However, if species B becomes extinct (*p* = 1 in panel b), its terminal branch is lost and species A becomes the sole representative of their shared internal branch, which then becomes an extension of species A’s terminal branch (blue branches in b). When the extinction risk of species B is between 0 and 1, as when *p* = 0.5 in panel (c), this represents a 50% probability that species B becomes extinct before a certain point in the future. There is therefore a 50% chance that species A, when its extinction risk is not considered, will become the sole representative of the shared internal branch between species A and B, and thus the ED2 of species A is now its terminal branch length (solid blue branch, panel c) plus 50% of the shared branch’s length (dashed blue and green branch, panel c).

From a technical perspective, ED2 is conceptually a special case of ‘expected distinctiveness’ [51] and identical to ‘Heightened Evolutionary Distinctiveness’ (HED) [40], expressed for species *i* as follows:

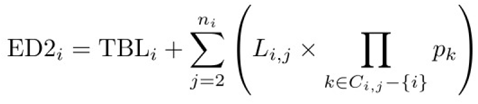

where ED2*_i_* comprises TBL*_i_* (terminal branch length of species *i*) plus another component. This is intuitive as the phylogenetic diversity captured by the terminal branch length is not shared with any other species whilst the other component captures shared contributions. The same is true of original ED, but here we write it explicitly. The set *C_i,j_* represents all species descended from the corresponding branch with length *L_i,j_* (as defined earlier) and *p_k_* is the probability of extinction of species *k* [37]. ED2*_i_* is therefore species *i*’s expected unique phylogenetic diversity (terminal branch length) at some point in the future assuming all other species in the tree survive or perish as a function of their probabilities of extinction (product of all *p_k_*). When species *i* has few close relatives, all with high *p*, it is expected to contribute a large amount of unique phylogenetic diversity because it is likely that all its relatives will perish, while a species with many secure relatives is not expected to contribute much more than its current terminal branch length. Changes in *p* for other species on the same branch therefore change the ED2 score of species *i.* The formula considers the branch corresponding to *L_i,j_* in turn and calculates its probability of becoming part of the terminal branch for species *i* in the future due to the extinction of all other descendants.

As our formulation of ED2 is equivalent to HED, the product of ED2 and GE2 makes EDGE2 mathematically equivalent to the ‘Heightened EDGE’ (HEDGE) approach where the future of the species is made secure([40] theorem 4.1 (i)). However, given it is unrealistic that a species can ever be completely ‘secure’, we conceptually consider EDGE2 scores to represent the potential expected PD loss that might be averted to some extent by conservation action on a single species [37]. As the success of conservation actions varies, one future change to the EDGE2 metric could incorporate to what extent different actions will impact the extinction risk of a species (e.g. downlisting by one Red List category), and thus GE2 could be weighted to reflect changes in extinction risk that assume different outcomes following conservation action [37,43].

#### Terminal versus internal branches

One consequence of ED2’s explicit incorporation of PD complementarity is the interaction between phylogenetic structure and extinction risk, through the multiplication of branch lengths and probabilities of extinction. This interaction naturally down-weights the contribution of deeper phylogenetic branches to the ED2 scores of species; the more descendant species there are, the smaller the probability that a branch will be lost. This is not necessarily a concern: most species-based PD metrics are heavily weighted by the branches closest to the tips [67,68], and terminal branches have long been recognised as a valuable measure of distinctiveness [2,8,66]. However, a key area of consensus at the workshop was that the contribution of internal branches to biodiversity should be valued in any EDGE framework.

To highlight the relative contributions of terminal and internal branches to ED2 scores, we explicitly decomposed ED2 into two components: (1) the terminal branch length, which represents the current PD exclusive to that species, and therefore its current distinctiveness [66] (Fig 2a); and (2) the cumulative contribution of all internal branches, accounting for PD complementarity, which represents the additional PD for which the species is expected to be responsible into the future given the current extinction risk of its relatives (Fig 2b, c). This facilitates the identification of species that are expected to be responsible for a large amount of PD from internal branches, indicating that they are part of highly distinctive clades with widespread elevated extinction risk. The decomposition of ED2 scores also permits terminal branch lengths—which are highly correlated with original ED [68]—to be used as a minimum estimate of ED2 when deeper phylogenetic structure is highly uncertain.

#### ED2 uncertainty

Since the inception of the EDGE metric, it has been the philosophy to incorporate all described species to ensure all species of conservation concern are included where possible, particularly those species from old lineages with few, if any, closely related species [19,22,24,25]. The original EDGE approach had a set of rules for resolving uncertainty in ED scores arising from lack of phylogenetic structure (polytomies) in poorly-known regions of the phylogeny—based on assumptions regarding net diversification rates—and also from the omission of species from the phylogeny, based on the observed ED scores of their closest relatives [19].

It is increasingly common for fully-resolved phylogenetic trees to be generated that include all described species of large clades (at that time), including those species for which we have no genetic data [28,69–74]. Such synthetic phylogenetic trees often utilise taxonomic information and established phylogenetic imputation methods (e.g. [75]) to generate what are essentially Bayesian posterior distributions of imputed phylogenetic trees [76]. This approach is inherently uncertain (more so than other phylogenetic hypotheses), particularly in regard to the phylogenetic structure of regions of the phylogenies with large proportions of imputed species, and this uncertainty must be reflected in the calculation of ED2 scores and the subsequent identification of conservation priorities [22,46]. ED2 is therefore calculated across a large distribution of imputed phylogenetic trees (100 < n ≤ 1000 to capture phylogenetic uncertainty [26,46]) that comprise all described species for the clade being assessed. This generates a Bayesian posterior distribution of scores from which we can determine the robustness of priority EDGE2 species using established criteria [46] (see ‘EDGE2 framework’ section, below).

#### EDGE2 framework

Both ED2 and GE2 scores for each species are not point estimates but a distribution generated by drawing many times from underlying distributions to account for uncertainty in phylogenetic relationships and extinction risks. The final EDGE2 score for a species is therefore a measure of the centre of a distribution (the median) of EDGE2 scores. Likewise, the final ED2 and GE2 values for a species are also the medians from their respective sampled distributions.

In the original EDGE framework, a priority ‘EDGE Species’ was one with an above-median ED score, relative to all species in the clade, which was also threatened (VU, EN or CR on the Red List) [19,22,24]. EDGE2 follows a similar philosophy: priority ‘EDGE2 Species’ must be noteworthy both in EDGE2 score and risk of extinction (GE2). We thus maintain the requirement for EDGE2 species to be in a threatened Red List category, but also include Extinct in the Wild species in this designation, given their potential for recovery in the wild. We also require the species to score above the median EDGE2 score for all species in the clade across at least 95% of the distribution of scores. This means we are 95% certain the species has an above median EDGE2 score after incorporating uncertainty in both extinction risk and phylogeny. Fig 3 summarises how we can use the Red List categories and EDGE2 scores to create lists for practical conservation prioritisation.

**Fig 3:**
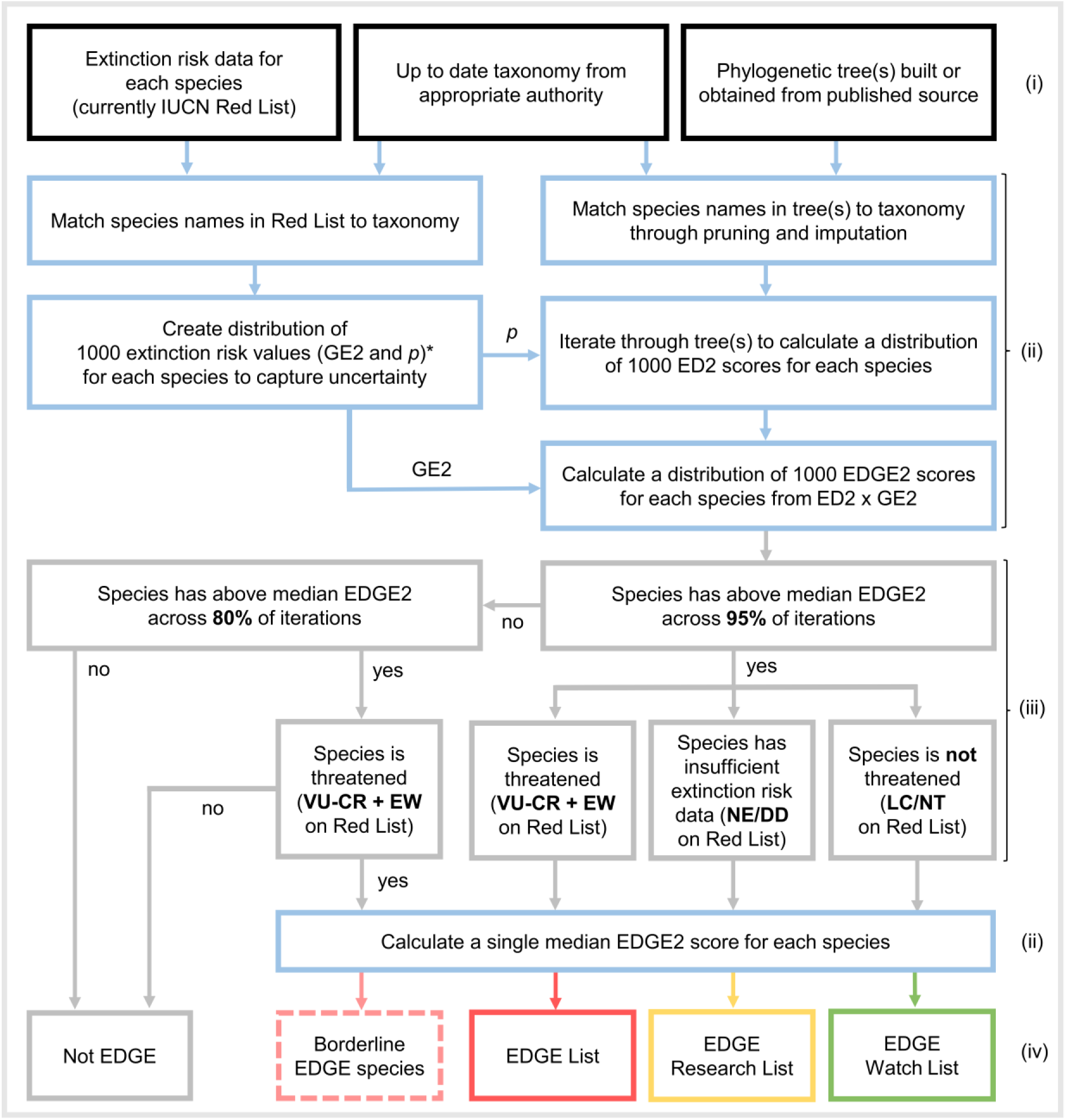
The EDGE2 framework. The workflow, from data input to final output, for producing a comprehensive EDGE2 prioritisation for a set of species. Stages from top (beginning) to bottom (end): required data (i – input data); data processing stages (ii – blue); selection criteria to be applied (iii – grey); and outputs (iv). \**p* and GE2 are assigned equivalent values in this study but may differ given increased understanding of extinction risk, the likely effects of future conservation work, on and its distribution across the Tree of Life (see ‘GE2’ section).

We here define an EDGE List as comprising all EDGE2 Species (those with EDGE2 values above the median across 95% of iterations and threatened with extinction; Fig 3). We also set criteria to highlight ‘borderline EDGE Species’; these are species that are threatened with extinction whose certainty around their EDGE score is still relatively high (e.g. above median EDGE2 >80% of the time) but not enough to meet our threshold of 95% certainty. For taxonomic groups with high phylogenetic uncertainty, these ‘borderline’ species would be included in the EDGE List with a relaxation of the certainty threshold (e.g. from 95% to 80%).

We also propose an ‘EDGE Research List’ for species that are credibly above median EDGE2 but are currently either NE or listed assessed as DD on the IUCN Red List (Fig 3). Given that the GE2 values for DD or NE species under EDGE2 are drawn from all possible values for data-sufficient categories, with a median approximately equivalent to that of VU species [45,64], any EDGE Research species are sufficiently evolutionarily distinct that they would become EDGE species if they were eventually to be assessed (or re-assessed for DD species) as VU or above on the IUCN Red List. We define an ‘EDGE Watch List’ for LC and NT species that rank above median EDGE2 95% of the time despite their low extinction risk, and are therefore responsible for securing large amounts of imperiled PD (Fig 3).

## Results

### EDGE2 for the world’s mammals

We calculated 1,000 values of GE2, ED2 and EDGE2 for 6,253 mammal species, 4,847 of which could be linked to IUCN Red List assessments with a data-sufficient Red List category. GE2 values ranged from 0.00176 (minimum for LC) to 0.999962 (maximum for Critically Endangered). Median ED2 scores across the 1,000 trees ranged from a minimum of 0.08 MY (*Uroderma bakeri,* Baker’s Tent-making Bat) to a maximum of 77 MY (*Orycteropus afer,* Aardvark), with a median ED2 of 2 MY (Fig 4b). The Large-headed Capuchin (*Sapajus macrocephalus*) had the lowest EDGE2 score (0.01 MY), and the species with the greatest median EDGE2 score (25 MY of avertable expected PD loss) was the Mountain Pygmy Possum (*Burramys parvus;* Table 1). The median EDGE2 score for mammals was 0.2 MY of avertable expected PD loss (Fig 4c).

**Fig 4:**
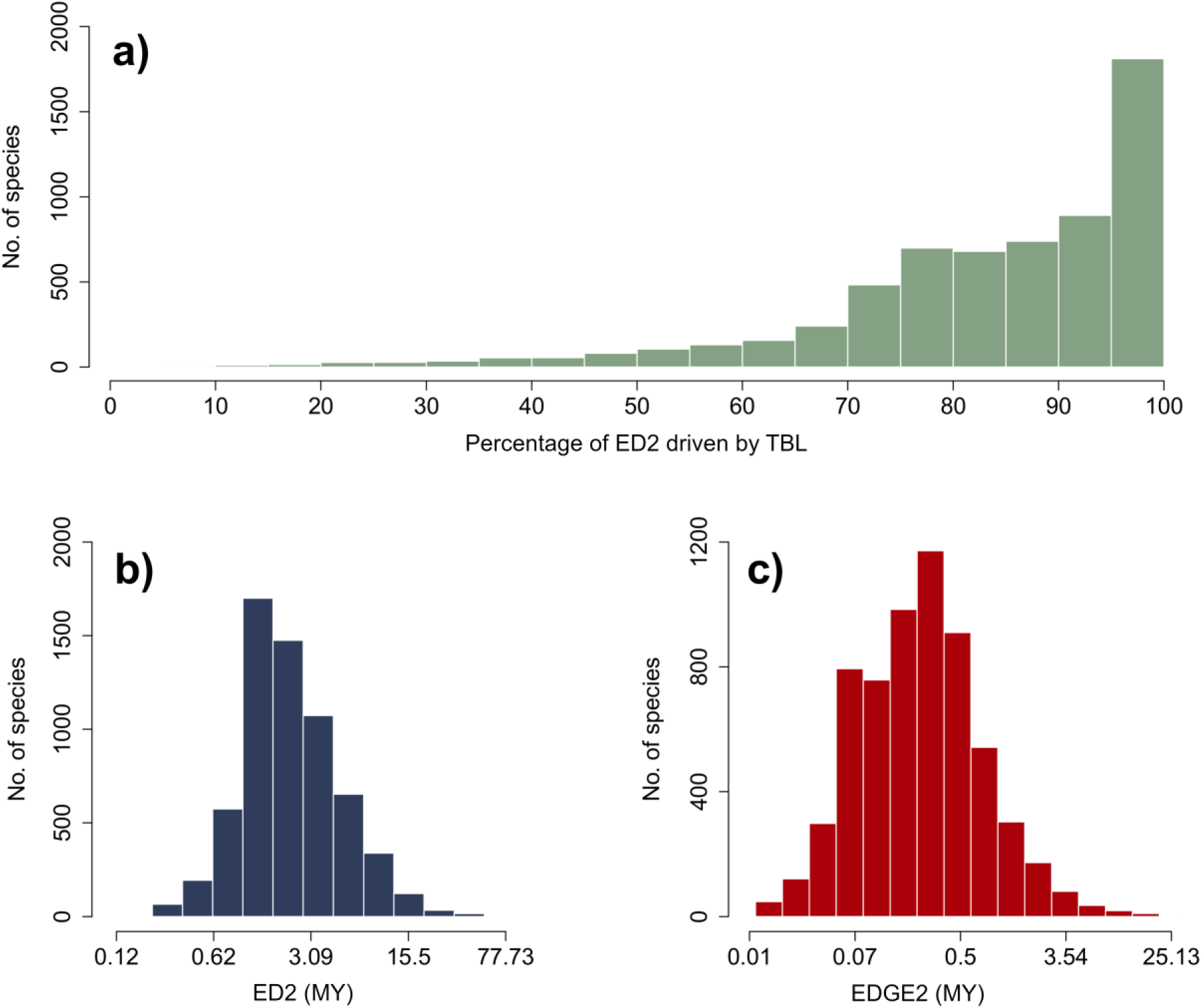
Distributions of ED2 and EDGE2 scores for mammals. The distribution of (a) the percentage of species’ ED2 scores contributed by their terminal branch lengths (TBL) alone; (b) ED2 scores; and (c) EDGE2 scores, for all mammals. Panels b and c are presented on a log-scale along the horizontal axis.

**Table 1:**
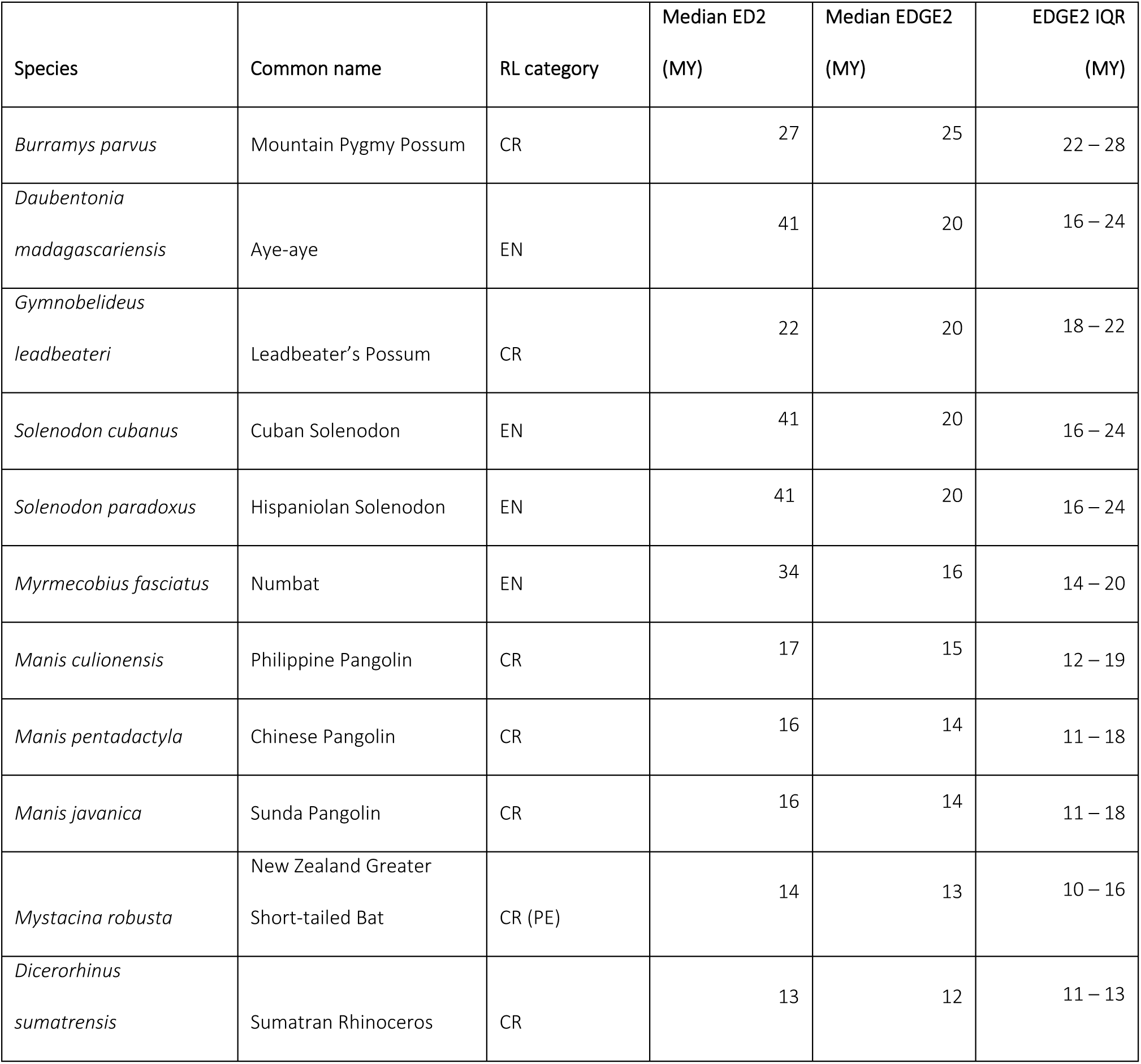
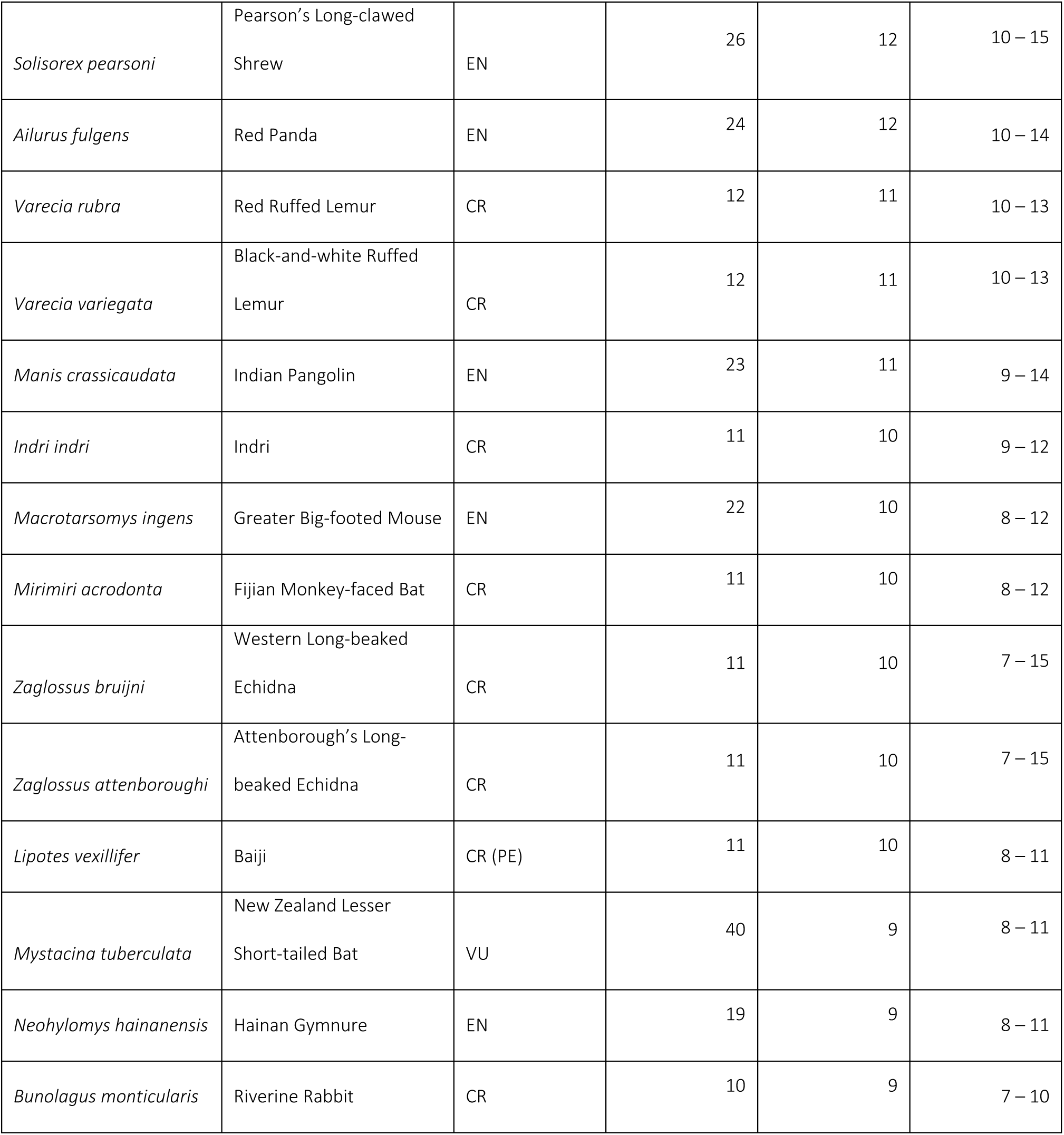
Top 25 EDGE2 Mammal species. The scientific name, common name, Red List category, median ED2, median EDGE2, and interquartile range of EDGE2 for the 25 highest-ranking EDGE2 Species. ED2 and EDGE2 scores calculated across a distribution of 1,000 phylogenetic trees. The Full EDGE List is given in the S1 File. All ED2 and EDGE2 scores are given to the nearest million years to reflect the degree of accuracy appropriate for these data given the uncertainty in phylogenetic and extinction risk estimates.

As expected (since their unique phylogenetic diversity is at risk of being lost), threatened mammal species account for almost 50% of all avertable expected PD loss (1,744 MY) despite accounting for only 22% of all mammal species diversity [77]. We identified 645 ‘EDGE2 Species’ (threatened species having above-median EDGE2 for 95% of iterations and VU, EN, CR, EW), which together account for 81% of the avertable expected PD loss of all threatened species (1,458 MY). Safeguarding the 100 highest-ranking threatened species (1.6% of mammal species) under EDGE2 would secure 719 MY of PD that would otherwise be lost, which is 41% of the total avertable expected PD loss for threatened mammals and 23% of avertable expected PD loss for all mammals (S1 Fig).

We identified 15 NE or DD mammal species that are above median EDGE at least 95% of the time (EDGE Research List; S1 File). Together these 15 species represented 44 MY of avertable expected PD loss. Six of the 15 species would be top 100 EDGE2 mammals if they were assessed as VU or higher on the Red List, and the highest-ranking species amongst these include the poorly-known Long-eared Gymnure (*Hylomys megalotis*; median ED2 = 34 MY) and Owl’s Spiny Rat (*Carterodon sulcidens*; median ED2 = 17 MY).

The EDGE Watch List comprises 82 species of LC (14 spp.) or NT (68 spp.) mammals that have above median EDGE2 across 95% or more iterations (S1 File). Together these species are expected to contribute 1,045 MY of PD, and the list includes highly evolutionarily distinct species such as the Aardvark (ED2 = 78 MY), Pin-tailed Treeshrew (*Ptilocercus lowii*; ED2 = 52 MY) and Duck-billed Platypus (*Ornithorhynchus anatinus*; ED2 = 51 MY).

### Comparing measures of ED

ED2 is strongly positively correlated with terminal branch length (TBL; Pearson’s product-moment: r = 0.95, df = 6251, p < 0.0001; S2 Fig). The median proportion of ED2 contributed by the TBL of the species is 87.2% (range: 4.8% - 99.9%; Fig 4a), and this proportion is moderately positively correlated with TBL alone (r = 0.326, df = 6251, p < 0.0001). ED2 is strongly positively correlated with original ED values calculated following Isaac et al. [19] (hereafter ED1; r = 0.829, df = 6251, p < 0.0001; S2 Fig), though ED2 scores are significantly smaller than those of ED1 (Wilcoxon signed rank: V = 5.58 x10^4^, df = 6251, p < 0.0001). This is due to the greater contribution of internal branches to ED1 scores and the fact that ED1 scores sum to PD whilst ED2 scores sum to a smaller figure (approximating total expected PD loss for the clade). The proportion of ED2 contributed by TBL is strongly negatively correlated with the proportional change in ED1 and ED2 ranks (r = -0.886, df = 6251, p < 0.0001). Sixty-three of the 100-highest ranking species under ED2 are also in the 100 highest ranks for ED1. Change in ranking from ED1 to ED2 is moderately positively correlated with the proportion of ED2 contributed by TBL (Spearman’s rank: ρ = 0.296, df = 6251, p < 0.0001), again reflecting the greater weighting towards terminal branches under ED2.

### Comparing EDGE priorities

EDGE2 ranks are strongly positively correlated with EDGE ranks calculated using the original Isaac et al. [19] approach (hereafter EDGE1) for all extant and data-sufficient mammals (ρ = 0.916, df = 4845, p < 0.0001; Fig 5; S3 Fig, see S4 Fig for comparison of raw EDGE values). Of the 645 EDGE2 Species identified here, 357 (60%) are also identified as priorities under EDGE1 Species criteria (VU, EN, CR and having above median ED1) when calculated using the same phylogenetic and extinction risk data. Amongst EDGE2 Species, which are the set of species identified as EDGE2 conservation priorities, higher priority species have smaller changes in their ranking from EDGE1 to EDGE2 compared with lower priority species (i.e. high-ranking EDGE2 priorities are more stable, having smaller rank changes between EDGE metrics; ρ = 0.378, df = 626, p < 0.001).

**Fig 5:**
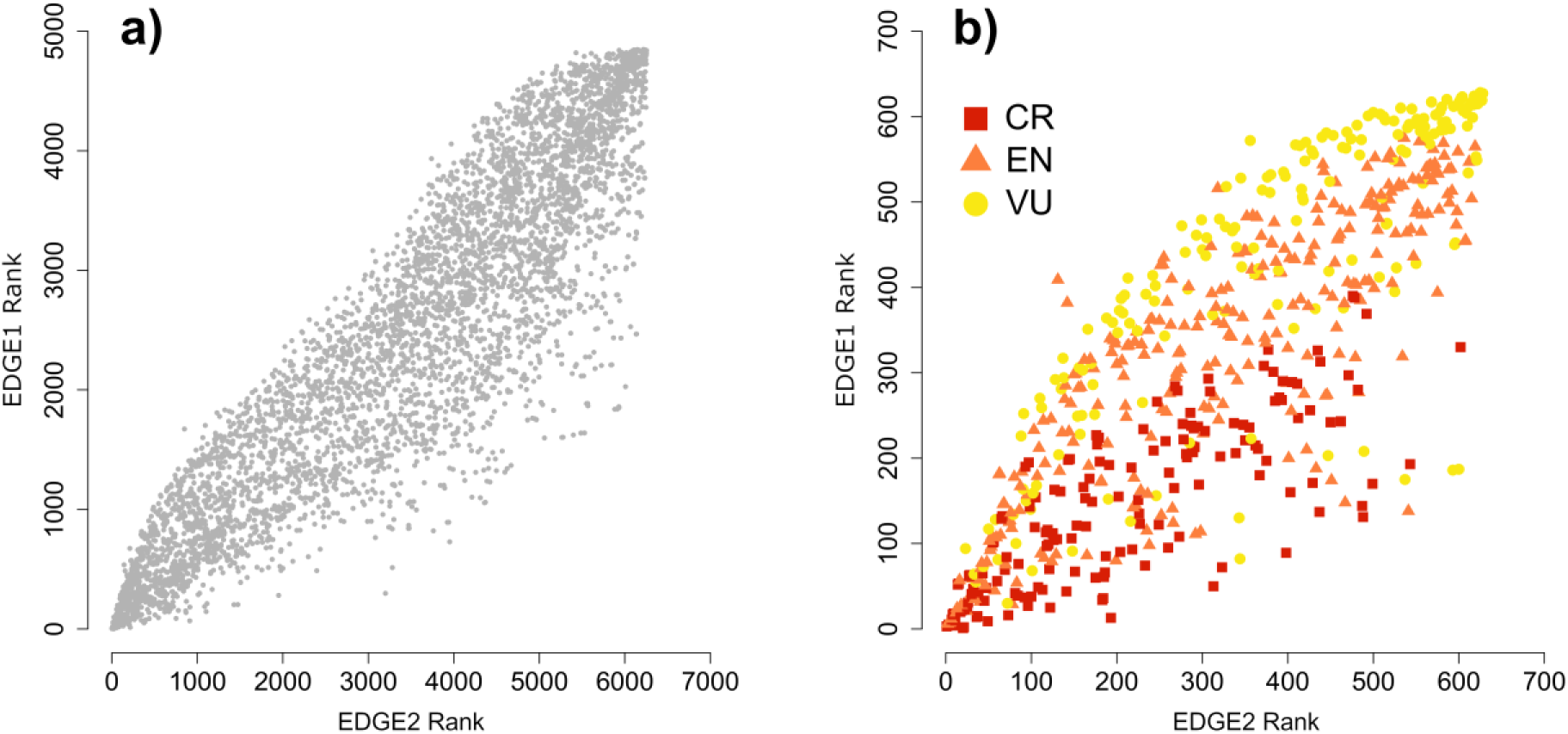
Relationship between EDGE1 and EDGE2 rankings. The EDGE2 ranks (x-axis) and EDGE1 ranks (y-axis) for (a) all mammals with data-sufficient and extant IUCN Red List assessments; and (b) all EDGE2 Species (above median EDGE2 for 95% of iterations and threatened on IUCN Red List; see ‘EDGE2 Framework’). Relationship between raw EDGE1 and EDGE2 scores in S4 Fig.

Sixty-seven of the 100 highest-ranking threatened species under EDGE2 are also in the top 100 ranking species under EDGE1. A further 30 species from the top 100 EDGE2 ranks are outside the top 100 EDGE1 ranks, but still qualify as priority EDGE Species under the original criteria. The three species not considered as EDGE Species under the original criteria are *Habromys lepturus*, *H. ixtlani*, and *Simias concolor*; they have ED1 scores below the median but EDGE2 scores above median. All of the top 100 EDGE Species identified using the EDGE1 framework are captured in the 645 EDGE2 Species, and 97 of the top 100 EDGE2 Species are captured in the 529 EDGE1 Species.

## Discussion

Here we have outlined EDGE2, an improved framework for identifying and prioritising species that are Evolutionarily Distinct and Globally Endangered (EDGE Species) for conservation action. Our updated EDGE2 metric builds on the original EDGE approach in three important ways. First, EDGE2 enables the use of probabilities of extinction—or other extinction risk weightings between 0 and 1—to represent the extinction risk of species, rather than the simple IUCN Red List Index equal-steps weighting of original ‘GE’. This makes EDGE2 more flexible, and thus capable of making use of future sources of, and improvements in the estimation of, extinction risk(e.g. [45]). Second, EDGE2 takes into account the extinction risk of closely-related species when calculating ED scores to better reflect the contribution a species is likely to make to the phylogenetic diversity of a group in the future (‘PD complementarity’) [66]. This bridges the gap between species-list approaches where each species is scored independently without the context of conservation or accounting for the state of other species (e.g. original EDGE lists), and set-selection approaches that produce sets of species as their output and do not aim to assign scores to individual species at all (e.g. [2,41]), making them complex to implement. Third, EDGE2 incorporates uncertainty around both phylogeny and extinction risk and utilises established methods to impute both the phylogenetic and extinction risk components for species lacking one or both to maximise the inclusivity of the metric. This allows us to cautiously calculate EDGE2 lists for more species for which there are uncertain data, broadening the scope and applicability of phylogenetic information in conservation.

### EDGE2 in theory

Conservation is a crisis discipline, evolving rapidly in both theory and practice, and requires decisions to be made quickly in the absence of complete information [78]. As such, conservation biologists strive to use the best information currently available, and the modular nature of EDGE2 facilitates the incorporation of improved data without rethinking the protocol in its entirety. For example, one limiting factor of the original EDGE metric is its reliance on available IUCN Red List assessments, and the mapping of Red List extinction risk categories to an existing index, to provide the ‘GE’ component of the metric. Our change under EDGE2 to a GE ‘module’ that weights extinction risk between 0 and 1 allows us to leverage Red List categories as the current best source of extinction risk data, but also gives EDGE2 the flexibility to make use of extinction risk weightings from other sources, such as Population Viability Analyses (PVAs) and quantifications of extinction probabilities [79], should such values ever become available for entire clades. The modular EDGE2 framework utilises the best available information, explicitly accounting for its uncertainty as underlying Bayesian posterior distributions, to generate hopefully robust prioritisations (Fig 3). This approach allows EDGE2 to be adaptable as theory, data, and conditions on the ground evolve.

The comprehensive nature of the EDGE2 framework is underpinned by the incorporation of uncertainty and imputation for both the extinction risk and phylogenetic placement of species where appropriate. It allows us to make use of the uncertainty generated by developing the comprehensive assessment of a clade to not only identify robust priorities for conservation action, but also to identify distinctive species lacking adequate extinction risk data for further research (EDGE Research Lists). The identification of such species has been proposed using various approaches in the past [25,48,80], but here they form a coherent part of the overall EDGE2 framework.

### EDGE2 in practice

We applied the new EDGE2 framework to the world’s mammals to explore the real-world implications of the new prioritisation. Mammals provided an ideal model with which to test the various modules of the EDGE2 framework: they have a recently-published distribution of phylogenetic trees that is virtually complete but requires the imputation of a small proportion of species, and the group is almost comprehensively assessed on the IUCN Red List. However, even for a group as well researched as mammals, 14% of species are Data Deficient, while several hundred recently described or split species remain to be assessed. This allowed us to generate a rigorous prioritisation whilst exploring the impacts of phylogenetic and extinction risk imputation on real-world results.

Overall, EDGE2 produces similar priorities to the EDGE1 approach applied to this new dataset, rather than wholesale changes. ED2 scores and ED1 scores are closely related, particularly when a large proportion of both ED scores is contributed by the terminal branches, for example in species that share their most recent ancestral internal branch with multiple species (i.e. species that are early-diverging in highly imbalanced lineages). This is best illustrated by the Aardvark, which is the highest-ranking ED species under both ED1 and ED2. The Aardvark has similar scores under ED1 (78.2 MY) and ED2 (77.7 MY), due to more than 99% of both ED scores being contributed by the terminal branch. The Aardvark shares its first ancestral internal branch with >70 spp. of sengis or elephant shrews (Macroscelidea) and tenrecs and golden moles (Afrosoricida), and thus receives a trivial proportion of its ED from this first internal branch under either metric. When ED scores are driven less by the terminal branch and more by the internal branches, the two ED measures become less similar, meaning that, under EDGE2, species with more close relatives that might all share a long ancestral branch are viewed as less of a priority than they are under EDGE1. This is a desirable feature of the new metric, as the greater number of extant descendant species, the less probable the loss of an internal branch becomes.

The new highest-ranking EDGE mammal, the Mountain Pygmy Possum, is the third-ranked species under the EDGE1 metric, behind the long-beaked echidnas *Zaglossus bruijni* and *Z. attenboroughi*. However, under EDGE2 the echidnas drop to 19^th^ and 20^th^, respectively. This is due to the high ED1, and therefore EDGE1, ranking of the echidnas being driven by their long, shared internal branches, which contribute more than 85% of their ED1 scores. However, with the incorporation of PD complementarity [40,66], the shared internal branches of the echidnas now contribute less than 50% of their ED2 scores, specifically because that internal branch is somewhat protected by the Near Threatened Duck-billed Platypus and the Least Concern Short-beaked Echidna (*Tachyglossus aculeatus*). Conversely, the Mountain Pygmy Possum receives >90% of its ED score from its terminal branch under both ED1 and ED2, and is therefore less impacted by the change in metric.

The explicit incorporation of PD complementarity into EDGE-like metrics has long been advocated for [40,41,43]. EDGE2 not only incorporates this complementarity to inform the identification of conservation priorities, but clarifies how this is done by partitioning the ED2 scores of species into their unique current contribution to global PD (terminal branch) and their additional expected future contribution due to the possible loss of related species (internal branches; Fig 1). This allows the identification of species whose ED2 scores are driven solely by their terminal branch distinctiveness (such as the Mountain Pygmy Possum and Aardvark) and species whose scores are also driven by their heightened responsibility for internal branches of the tree (as with the long-beaked echidnas). The conservation of priority species under both scenarios is crucial if we wish to avoid particularly large losses of PD [81].

In both the original EDGE1 and EDGE2 metric, some clades emerge as consistent priorities. For example, pangolins (Pholidota), of which four of the eight species are in the 25 highest-ranking EDGE2 Species (Table 1), and all eight are priority EDGE2 Species. Pangolins are also one of only two orders (along with Sirenia - manatees and dugongs) in which all species are priority EDGE2 Species. Though one-third of the 100 highest-ranking species (the original cut-off for highest priority EDGE Species [19]) do not overlap between the EDGE1 and EDGE2 metrics, both metrics perform well in capturing the highest priority species of the other. Indeed, EDGE2 Species of higher priority are less prone to large changes in rank between the two metrics than lower ranking priority species. We interpret this to mean that the original EDGE metric, whilst lacking some of the conceptual advances now incorporated into EDGE2, performs relatively well at accurately identifying priority species.

Further, as the distribution of ED scores is skewed due to the presence of species isolated on long phylogenetic branches, when these species are also threatened (i.e., high GE), they are likely to be priorities irrespective of the prioritisation metric used. Indeed, despite the use of a different set of mammal phylogenies and GE values, of the 11 species shared between the top 25 HEDGE mammals of Robuchon et al. [82] and the top 25 EDGE2 mammals here, seven were monotypic genera and had terminal branch lengths in the top 5% of all mammals, and all 11 had ED2 scores in the top 5% for all mammals. However, the choice of GE values has been shown to affect priorities identified for both original EDGE and expected PD loss approaches such as HEDGE and EDGE2 [25,42,43]. Consequently, there is still relatively low congruence between the top 25 priority mammals identified here and those of Robuchon et al. [82], identified using different data and criteria, despite the metrics applied being conceptually equivalent. Thus, it is important to establish a coherent and comprehensive framework, such as EDGE2, to ensure consistency across taxa to reliably inform conservation efforts.

The EDGE Research List for mammals includes several highly distinctive species with limited conservation knowledge, including the Long-eared Gymnure, a primitive hedgehog which may have diverged from its closest living relatives more than 30 million years ago. The expansion of the EDGE framework to explicitly prioritise species lacking sufficient conservation knowledge for further research extends previous efforts to catalyse targeted extinction risk assessments for highly distinctive species potentially at risk of extinction [25,83]. EDGE Research Lists can inform ongoing efforts to improve our understanding of extinction risk for priority Data Deficient species, such as the IUCN Species Survival Commission’s EDGE internal grants [38] and the Mohamed bin Zayed Species Conservation Fund.

The EDGE Watch List is topped by non-threatened mammals that are highly evolutionarily, ecologically and morphologically distinct, including possibly the world’s smallest mammal, the Bumblebee Bat (*Craseonycteris thonglongyai*) [84], the smallest extant anteater, the arboreal and nocturnal Silky Anteater (*Cyclopes didactylus*) [85], and the chronic alcohol-consuming Pen-tailed Treeshrew [86]. Of the 82 species on the Watch List, 68 are already Near Threatened and therefore close to qualifying as threatened and thus as full EDGE species, highlighting the importance of recognising them in this way.

### Future directions

While our EDGE2 framework synthesises numerous advances in the field of phylogenetically-informed conservation prioritisation, it also provides a clear picture of what further research is now needed to improve the various modules comprising the framework. We selected reasonable weightings of extinction risk that reflect relative weightings of earlier species-based prioritisations [19,21] applied to a politically and biologically relevant time period (50 years). However, there are numerous ways to incorporate extinction risk weightings into the framework [42,43,53,79,83], and we do not claim our approach to be definitive. As our understanding of extinction risk improves, we certainly expect this component of EDGE2 to be revisited, with the ultimate goal of a mechanistic and biologically accurate calculation of globally threatened PD.

The incorporation of species lacking extinction risk data into global assessments of biodiversity loss is increasingly recognised as important [47,48,65,83,87–90]. Here our approach to the incorporation of species lacking adequate extinction risk information was designed to retain simplicity and applicability to groups about whose extinction risk we still know relatively little (e.g. ray-finned fish, invertebrates, plants, and fungi). However, there are other possible approaches to assigning GE2 scores to DD/NE species, for instance drawing them in proportion to the observed frequency of extinction risk categories of either the assessed members of the clade or species that transitioned from DD to a data-sufficient category, or using extinction risk correlates to inform the imputed extinction risk of species [48]. These options require further exploration in the context of the EDGE2 framework.

The capability of EDGE2 to generate comprehensive prioritisations for clades for which we lack comprehensive data—which, realistically, is all large clades of animals, plants, and fungi—provides the possibility for EDGE prioritisations to be developed for large regions of the Tree of Life, beyond those clades already assessed under the original EDGE framework (amphibians, birds, corals, reptiles, sharks and rays, gymnosperms). Continuing advances in open-sourced estimates of the Tree of Life [91] and their synthesis with extinction data [92], paired with the potential for increased IUCN Red Listing of large groups of species, means EDGE prioritisations for large portions of the Tree of Life are a realistic goal. Further, given that the vast majority of a species’ ED2 and EDGE2 scores are contributed by branches closest to the tips of the phylogenetic tree, EDGE2 scores can be considered comparable across various clades when calculated from separate phylogenetic trees (e.g. squamates, crocodilians and turtles), as the deep internal branches will contribute trivial amounts to ED2 scores due to the inclusion of PD complementarity [68]. This advance provides us with the option to generate and explore the use of EDGE Lists of species from disparate clades based on other thematic or taxonomic criteria (e.g. EDGE Mangroves, EDGE Pollinators, or Rankings of EDGE species listed on CITES).

### Conclusions

EDGE2 arrives on the eve of a major new framework for biodiversity conservation that the Parties to the Convention on Biological Diversity (CBD) will finalise in 2022. This vision will include a strident new call to turn the tide for species conservation, specifically to prevent further species extinctions, stabilize net extinction risk and to drive an increase in average population size. We believe the EDGE approach has a critical role to play in achieving this ambitious plan, especially in helping to prioritise investments in conservation efforts. However, the science behind the original EDGE metric has evolved considerably since the metric was originally proposed, and we saw the EDGE’s 10-year anniversary as an opportunity to revisit the EDGE methodology. By developing EDGE2 we hope to ensure that, as theory and data availability for conservation prioritisation continues to grow, more initiatives can implement conservation action underpinned by robust priority setting for another decade and beyond. Our new EDGE2 metric uses a modular, tiered approach to provide EDGE2 scores—and associated measures of uncertainty—for all species in a clade. One of the great strengths of the EDGE of Existence programme is its positive interactions with the wider scientific community, and our workshop and ongoing collaboration have allowed us to include that community in the EDGE2 process. We feel that EDGE2 is an example of how academic and applied conservation biologists can work together to guide effective priority-setting that has the best chance of preserving the biodiversity upon which humanity depends.

## Materials and methods

### Data

We applied the principles of EDGE2 to develop a revised global priority list for the world’s mammals. We selected a random sample of 1,000 phylogenetic trees from the recently-published [73] 10,000-tree ‘‘pseudo-posterior’ distribution of birth-death node-dated trees comprising 5,911 extant and extinct mammal species. Our taxonomy for mammals was taken from version 1.1 of The Mammal Diversity Database [93], from which we determined 6,253 extant valid mammal species as of 01/05/2020. We matched this taxonomy with both that of our distribution of phylogenetic trees and with IUCN Red List data for 5,853 mammal species (as of 11/05/2020; IUCN 2020). We removed all Extinct species from each tree to remove their influence on the ED2 and EDGE2 scores of closely-related taxa [22] and to more accurately estimate expected PD loss for extant species only, though for EDGE2 prioritisations their influence can also be removed by setting their extinction risk weighting to 1. To ensure EDGE2 priority setting for mammals was comprehensive, we then inserted all valid mammal species—according to our taxonomy—missing from the phylogenetic trees (421 spp.; 6.7% of mammals) to their respective genus (or family for species with no congeners in the tree) using the ‘*congeneric.impute*’ function in the R package ‘*pez*’ [94]. This approach approximates a previous method [76] for imputing missing species into a phylogeny. We generated a distribution of 1,000 trees comprising all extant mammal species at the time of analysis.

### EDGE2 for the world’s mammals

To generate a distribution of probabilities of extinction for each IUCN Red List category we fitted a quartic curve through the five median values for each Red List category (LC = 0.060625, NT = 0.12125, VU = 0.2425, EN = 0.485, CR + EW = 0.97) and bounded the resulting curve to return values between 0.0001 and 0.9999. We calculated ED2 and EDGE2 across our distribution of 1,000 trees and, where for each tree we selected new extinction risk weightings for each species at random from the distribution of GE2 scores tied to the species’ Red List category. This resulted in 1,000 ED2 and EDGE2 scores for all species. For currently NE and DD species we drew their GE2 scores from the entire distribution of GE2 values, which had a median of 0.232, comparable to that of the VU category (S5 Fig), and consistent with trait and range size-based predictions that indicate elevated extinction risk amongst DD species [64,65]. We then generated the EDGE Species List, as well as the Watch, Research and Borderline EDGE Lists for mammals (Fig 3). All ED2 and EDGE2 scores are given to the nearest million years to reflect the degree of accuracy appropriate for these data given the uncertainty in phylogenetic and extinction risk estimates. Functions to generate GE2 and EDGE2 scores are available online to speed up the generation of future EDGE2 lists (https://github.com/rgumbs/EDGE2/).

### Comparing measures of ED

The proportion of the ED2 score contributed by the TBL indicates the relative importance of the current distinctiveness of the species in question (large proportion of ED2 contributed by TBL would be a species with high ‘PD endemism’, *sensu* [66]) versus the heightened potential future responsibility of the species for internal branches due to the high probability that the other species descending internal branches will become extinct (large proportion of ED2 contributed by internal branches). We therefore calculated the terminal branch length (TBL) for each species in each tree and ran a correlation of ED2 and TBL to explore whether larger ED2 scores were associated with longer terminal branches or with the component corresponding to interior branches. To determine whether species with longer terminal branches receive a greater proportion of their ED2 from their TBL than those with shorter branches, we correlated TBL and the proportion of ED2 contributed by the TBL for each species.

We calculated original ED (hereafter ED1) scores across the 1,000 trees for all species and correlated the median ED1 scores for each species with their ED2 scores. We correlated ED2 ranks with ED1 ranks and calculated the percentage overlap between the highest 100 ranking species between the two. We also correlated the change in rank from ED1 to ED2 for each species with the proportion of ED2 contributed by TBL for each species to determine whether species with greater contributions from their terminal branches rank more highly under ED2 than ED1, given the latter accounts for the complementary contribution of close relatives to the persistence of internal branches.

### Comparing EDGE priorities

For comparison with the new EDGE2 approach, we calculated original EDGE (EDGE1) scores for all species with data-sufficient Red List categories (LC, NT, VU, EN, CR), and limited the EDGE2 comparison dataset to species assigned to the same Red List categories. We correlated EDGE2 ranks with EDGE1 ranks and calculated the percentage overlap between the highest 100 ranking species of the two.

Finally, we explored how the magnitude of change in EDGE rank from EDGE1 to EDGE2 was related to the ranking priority of the species to determine whether highly ranked EDGE2 species varied less in their rank change than lower-ranked (lesser priority) EDGE2 species.

One concern of using imputed phylogenetic trees, as we have here, is the impact of the imputed species on the accuracy of the ED2 and EDGE2 scores of the species for which we have molecular data [22,46], and the relative stability of EDGE2 scores for these imputed species. We conducted sensitivity analyses to explore the impact of imputing missing species at different stages of phylogenetic construction on EDGE2 scores (see S1 Text).

## Acknowledgments

We thank Aki Mimoto for uncovering the Shapley Value for phylogenetic trees, and Gavin Thomas, Klaas Hartmann, Tyler Kuhn, and David Redding for years of work and discussion. We thank Jonathan Baillie for his support of the EDGE2 workshop and the growth of ZSL’s EDGE of Existence programme. This research, through JR and WDP, is an output of the Georgina Mace Centre for the Living Planet at Imperial College London. Sadly, this manuscript was finalised after the passing of Georgina Mace. We are incredibly grateful for the contributions provided by Georgina during the EDGE2 workshop and early stages of this research, and supporting work on the EDGE concept from the very beginning.

## Supporting information

**S1 Text. Imputation analysis methods and results.**

**S1 File. EDGE2 scores for the world’s mammals.**

**S1 Fig. Proportion of EDGE2 captured by increasing numbers of species.** Cumulative percentage of total EDGE2 captured by increasing the number of species captured, from the mammal species with the highest EDGE2 score to the lowest.

**S2 Fig. The relationship between measures of evolutionary distinctiveness.** Relationship between median (a) terminal branch length (TBL) and ED2 scores; (b) ED1 and ED2 scores; and (c) TBL and ED1 scores, for all mammals.

**S3 Fig. The relationship between EDGE2 and EDGE1 ranks.** The relationship between EDGE2 and original EDGE ranks for (a) all assessed; (b) Least Concern; (c) Near Threatened; (d) Vulnerable; (e) Endangered; and (f) Critically Endangered mammal species.

**S4 Fig. The relationship between EDGE2 and EDGE1 scores for mammal species**. EDGE2 has been log-transformed for comparison with the log-like EDGE1 scores (see ‘The history of the EDGE metric’ section).

**S5 Fig. Mapping GE2 to extinction risk weightings.** Distribution of GE2 scores from which extinction risk weightings were selected for each Red List category, corresponding to their original GE1 scores in EDGE1. For GE1 scores, 0 = LC, 1 = NT, 2 = VU, 3 = EN, 4 = CR. DD and NE species were not given a GE1 score in the original EDGE1 metric.

## Notes

### Competing Interest Statement

The authors have declared no competing interest.

