## Supplementary material for "EDGE2: advancing the prioritisation of threatened evolutionary history for conservation action": S1 Text

### Imputation analyses

One concern of using imputed phylogenetic trees, as we have here, is the impact of the imputed species on the accuracy of the ED and EDGE scores of the species for which we have molecular data (Weedop et al. 2019). To explore this, we calculated EDGE2 rankings for all species in both the molecular-data-only (hereafter 'molecular-only tree') phylogenetic tree of Upham et al. (2019) and the larger tree containing species they imputed using taxonomy (hereafter 'taxonomy-imputed tree'). We correlated these rankings with one another for all species present in both trees, and also correlated both sets of rankings from the Upham et al. trees with those from our phylogenetic trees with the remaining 421 missing species imputed (hereafter 'fully-imputed tree'). To shed light on the impacts of our imputation on conservation priorities, we determined how many species in the top 100 EDGE2 ranks have i) molecular data, ii) were imputed by Upham et al. (2019), and iii) were imputed in our study. Finally, to determine whether the imputation of species increased variation in ED2 and EDGE2 scores, we calculated the variance in ED2 and EDGE2 scores for all species across the 1,000 trees and correlated these against the proportion of each genus that was included in the phylogenetic tree using genetic data.

The median EDGE2 rankings from our fully-imputed mammal phylogenetic trees are strongly correlated with those from the taxonomy-imputed tree ($\rho = 0.99$, df = 5401, $p < 0.0001$; Figure A). The EDGE2 rankings from our fully-imputed trees are strongly correlated with those from the molecular-only tree ($\rho = 0.79$, df = 4020, $p < 0.0001$), and the sets of EDGE2 rankings from the taxonomy-imputed and molecular-only trees are similarly correlated ($\rho = 0.80$, df = 4020, $p < 0.0001$; Figure A). Eighty-eight of the 100 highest ranking EDGE2 mammals were placed in the phylogenetic tree using molecular data, and the remaining 12 species were imputed based on taxonomy by Upham et al. (2019). Just six of the 633 EDGE2 Species (above median EDGE2 for 95% of iterations and VU, EN, CR) were imputed in this study.

Species imputed here have variance in both ED2 and EDGE2 scores comparable to those imputed by Upham et al. (2019), though both exhibit greater variance than species placed using molecular data (Figure B). Variance decreases as molecular coverage of a genus increases for both ED2 ($\rho = -0.394$, df = 6251, $p < 0.0001$) and EDGE2 scores ($\rho = -0.221$, df = 6251, $p < 0.0001$). This increased variance meant that of 421 species imputed here with EDGE2 scores above the median, just 7 (1.6%) were above median EDGE2 in 95% or more iterations, compared with 33.8% of species with molecular data. This increased variance limits the potential for species for which we lack adequate understanding of their evolutionary distinctiveness to dominate EDGE2 priority lists unless the species are part of particularly ancient and species-poor clades. The inclusion of otherwise missing species also serves to reduce the ED2 and EDGE2 scores in clades where a small proportion of described species are included, such as the genus *Dromiciops*, where two recently-described species were absent from the Upham et al. (2019) phylogenetic tree (Burgin et al. 2018), thus reducing the potential for species to be incorrectly identified as priority species due to overestimation of their evolutionary distinctiveness.

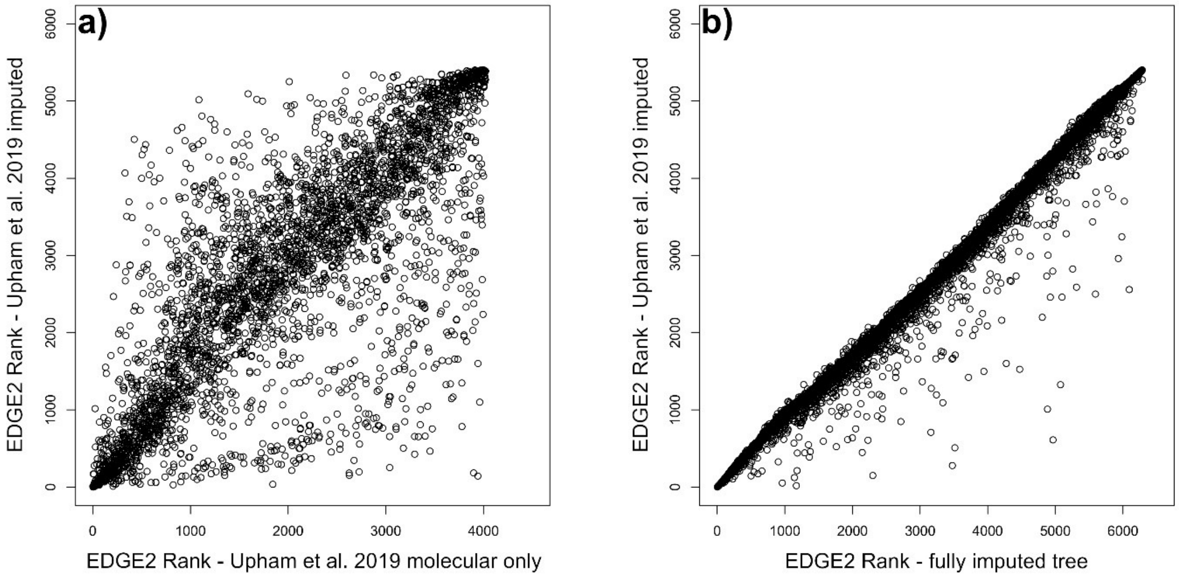


*Figure A: Relationship between the median EDGE2 ranks from (a) the taxonomy-imputed tree and molecular-only tree of Upham et al. (2019); and (b) the taxonomy-imputed tree of Upham et al. (2019) (y-axis) and the fully-imputed tree generated from this study (x-axis).*

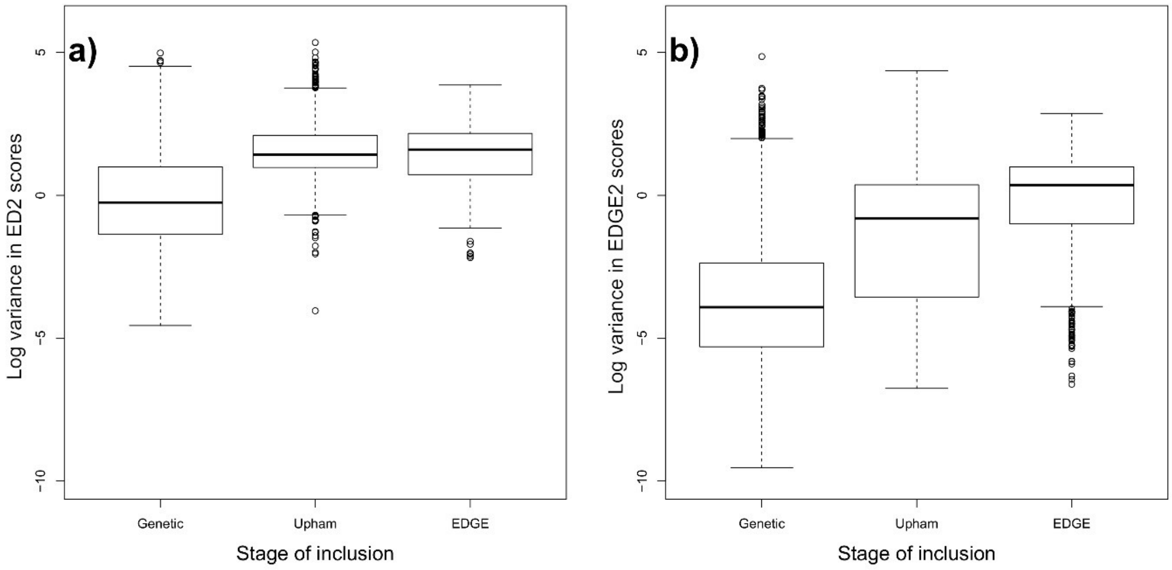


*Figure B: Variance in the (a) ED2 and (b) EDGE2 scores for species inserted into the phylogenetic tree using either genetic data, taxonomy by Upham et al. (2019), or taxonomy here, calculated across 1,000 phylogenetic trees.*
