## Supplementary figures and images for "EDGE2: advancing the prioritisation of threatened evolutionary history for conservation action"

### S1 Fig

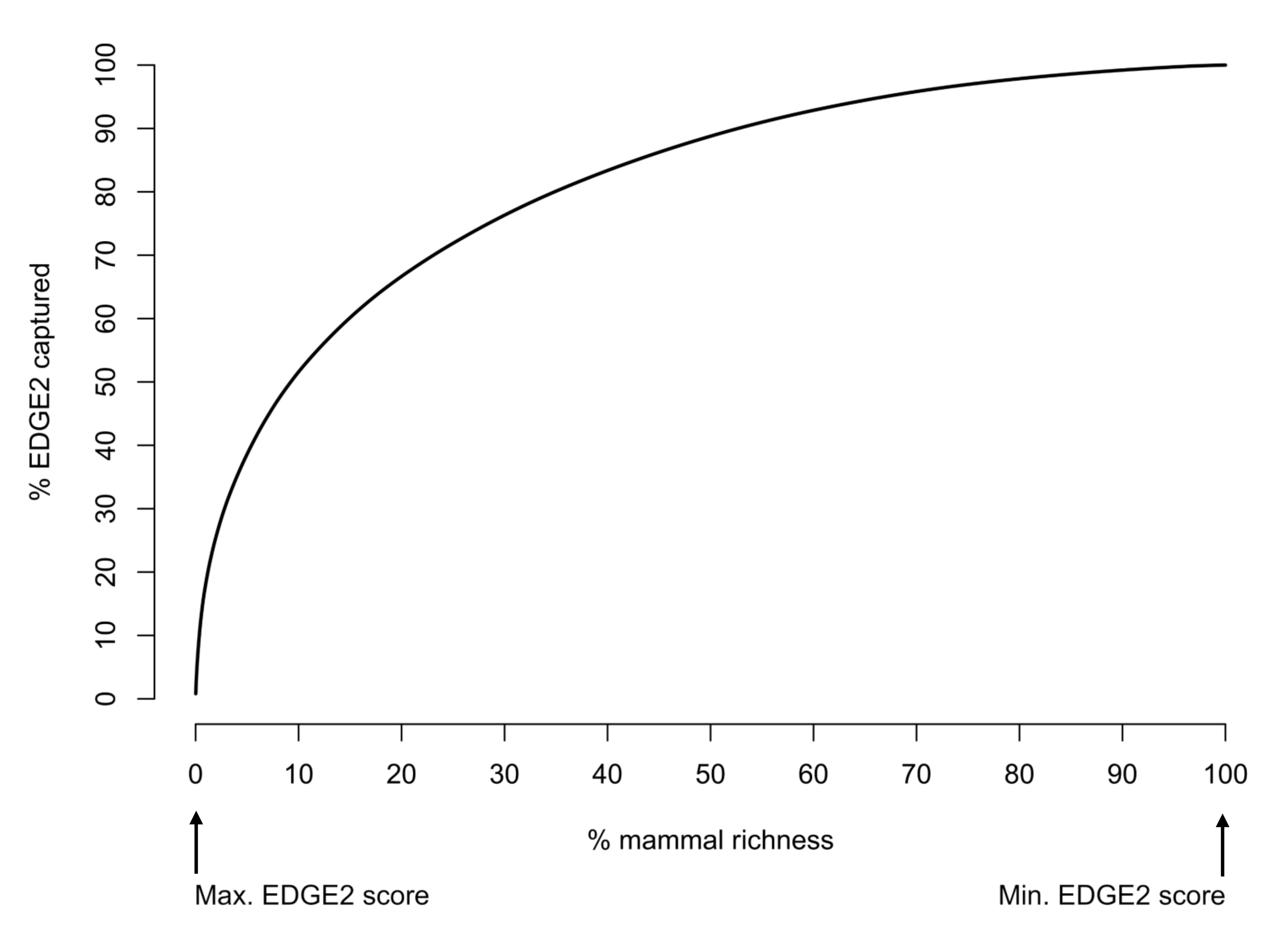

### S2 Fig

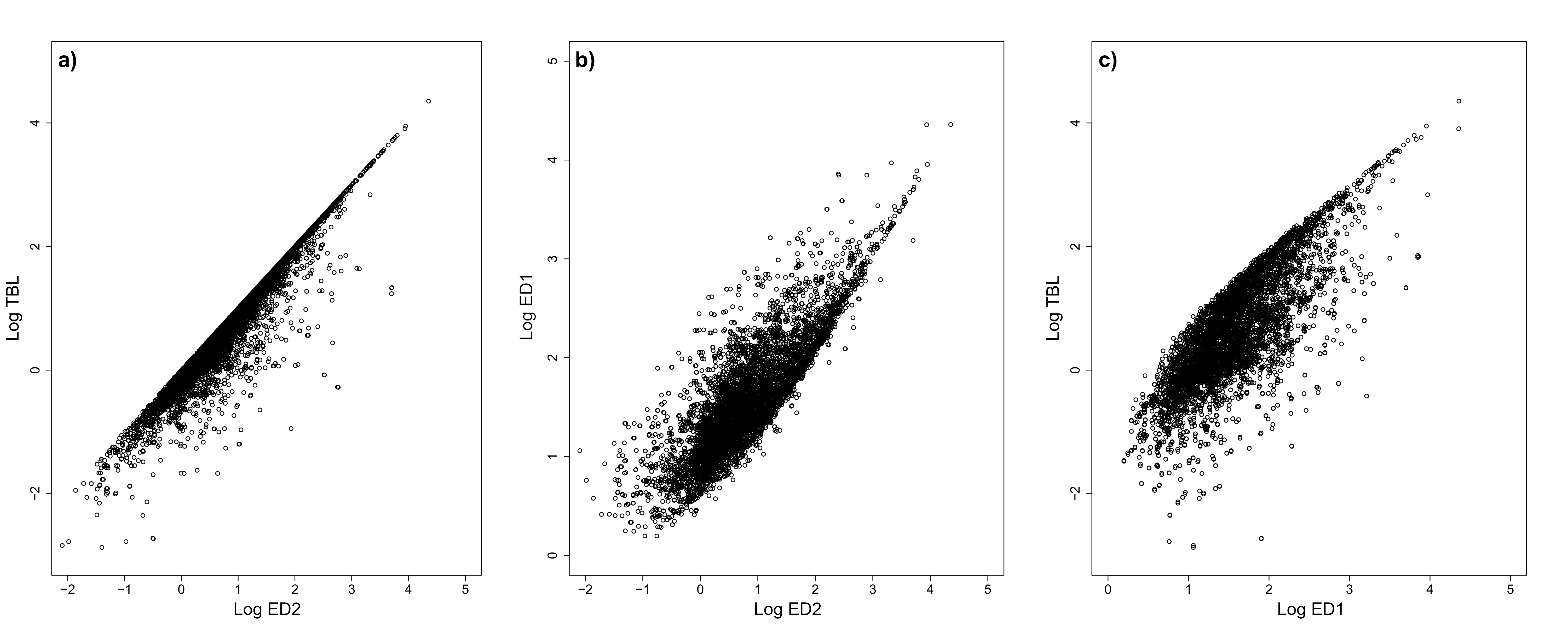

### S3 Fig

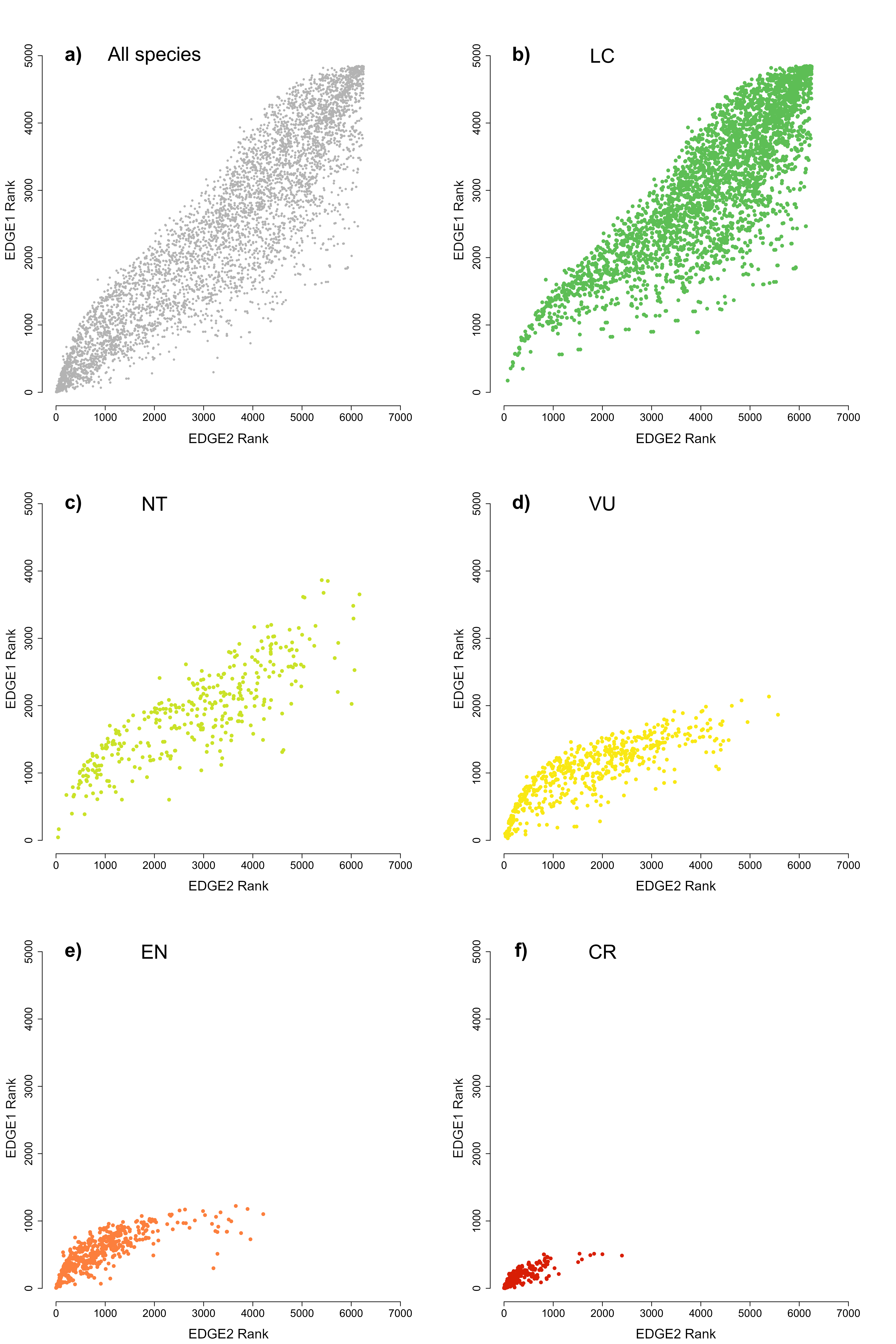

### S4 Fig

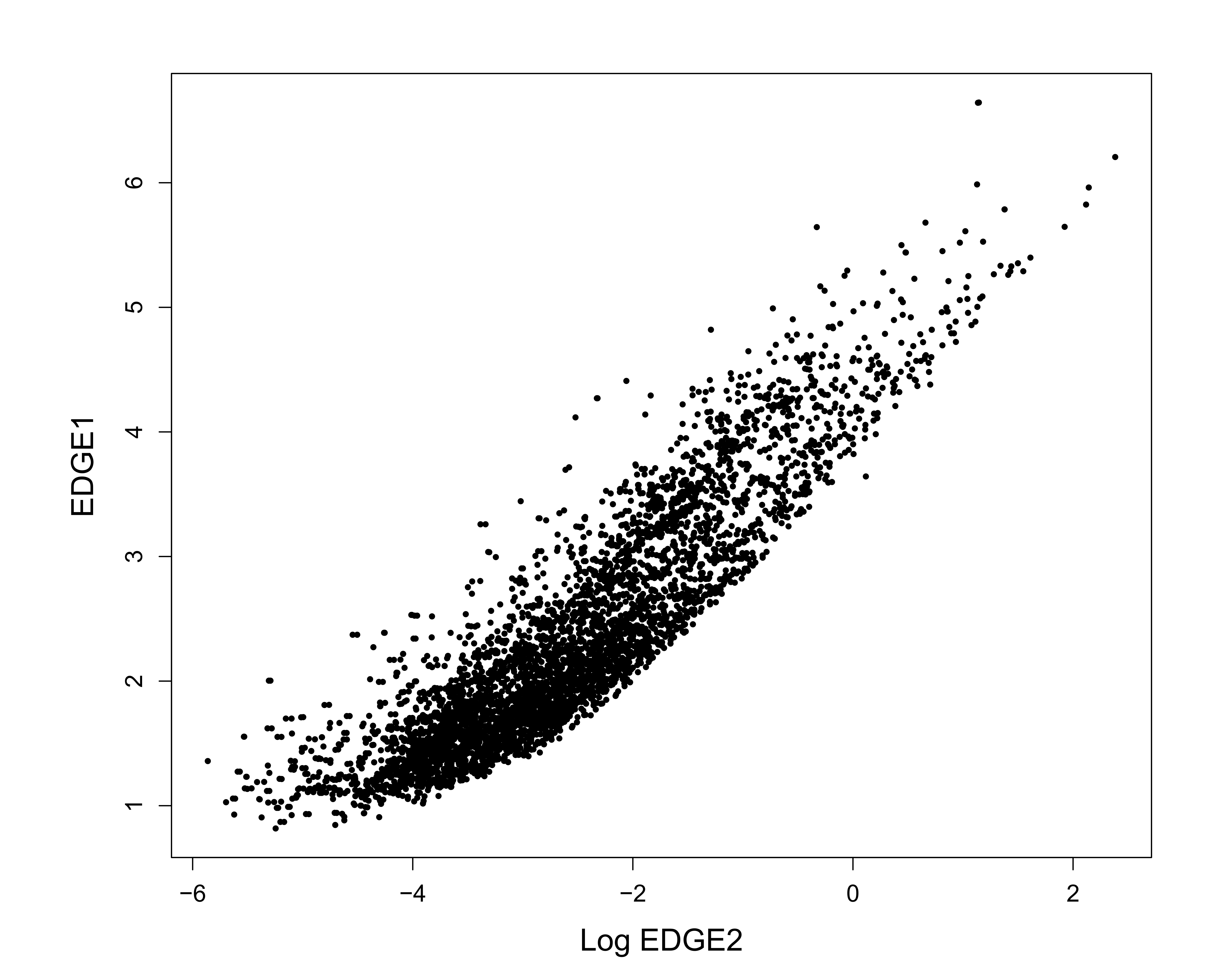

### S5 Fig

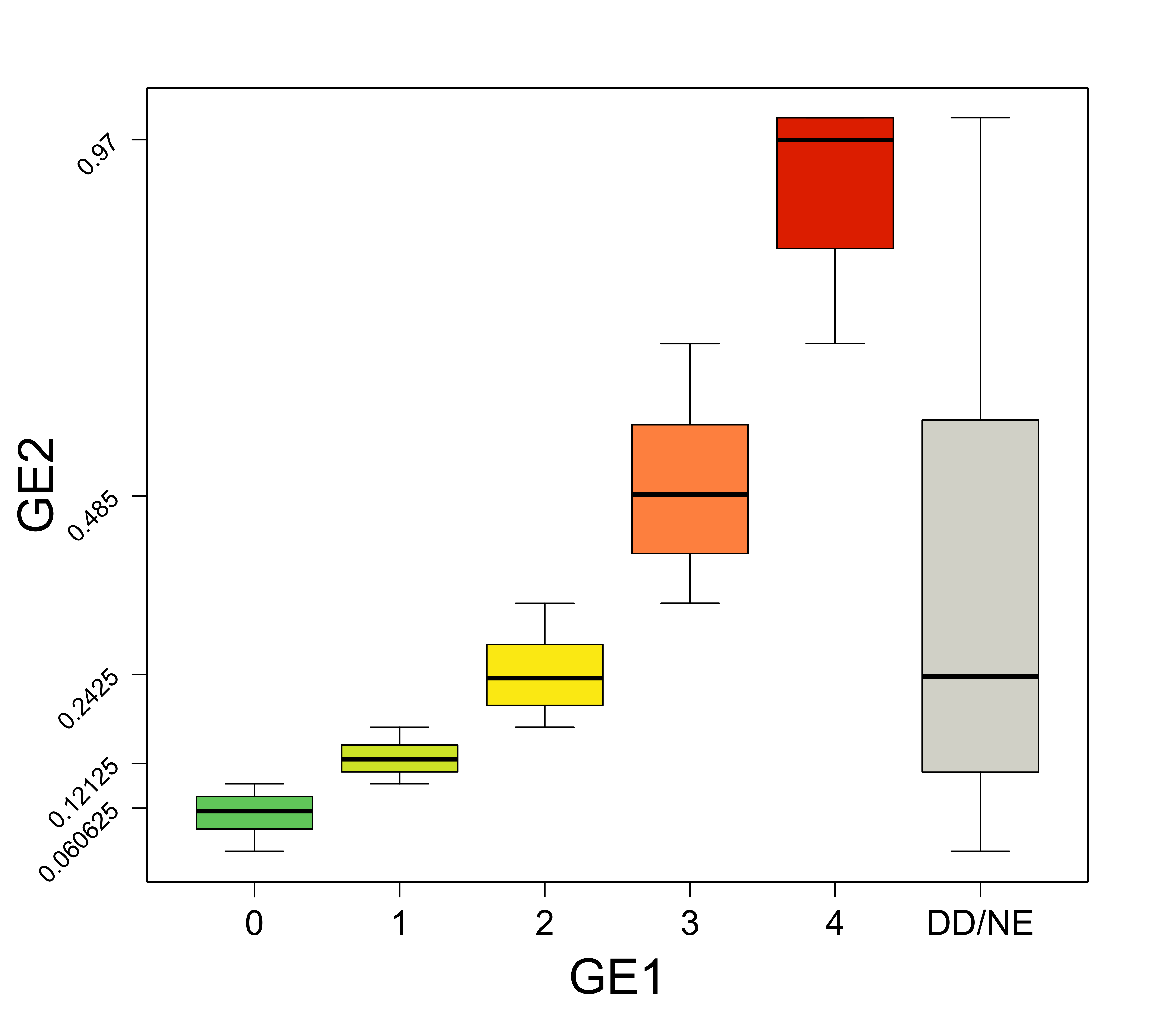
